# Functional characterization of cytochrome P450s associated with pyrethroid resistance in the olive fruit fly *Bactrocera oleae*

**DOI:** 10.1101/2022.12.07.519177

**Authors:** Anastasia Kampouraki, Dimitra Tsakireli, Venetia Koidou, Marianna Stavrakaki, Stavroula Kaili, Yannis Livadaras, Linda Grigoraki, Panagiotis Ioannidis, Emmanouil Roditakis, John Vontas

## Abstract

Resistance to pyrethroid insecticides has evolved in *Bactrocera oleae* populations in Greece, threatening the efficacy of control interventions based on this insecticide class. Here we report the collection of populations from Crete with resistance levels reaching up to 331-folds, compared to susceptible laboratory strains and show that pyrethroid resistance is substantially suppressed by the PBO synergist, suggesting the involvement of detoxification enzymes. To identify specific candidate genes implicated in resistance, we performed comparative transcriptomic analysis, between the pyrethroid resistant populations from Crete and the susceptible laboratory strains, using both whole bodies and Malpighian tubules. Several genes were found differentially transcribed between resistant and susceptible flies in each comparison, with P450s being among the most highly over-expressed detoxification genes in pyrethroid resistant populations. Four of the over-expressed P450s (*Cyp6A61, Cyp6G6, Cyp4P6* and *Cyp6G28*) were recombinantly expressed in *Escherichia coli* and *in vitro* metabolism assays revealed that CYP6A61 is capable of metabolizing alpha-cypermethrin, while CYP6G6, CYP4P6 and CYP6G28 are capable of metabolizing deltamethrin. No metabolism of neonicotinoid insecticides was recorded. We further silenced *CYP6G6 in vivo*, via RNAi, which led to a small, but significant increase in deltamethrin toxicity. The study provides valuable information towards the development of molecular diagnostics and evidence-based insecticide resistance management strategies.

## 1. Introduction

The olive fruit fly, *Bactrocera oleae*, is the major pest of olive orchards worldwide. This translates into enormous losses for the economy of regions that are based on olive oil production (Daane and Johnson, 2010). For example, according to Association of Cretan Olive Municipalities, in 2019 in Crete (Greece), the damage to olive cultivation by olive fruit flies exceeded 80,000,000 euros (sedik, 2019).

The control of *B. oleae* in Greece and the Mediterranean basin is largely based on the use of chemical insecticides. Other methods, such as mass trapping are also applied. Recent advances in the biotechnology-based SIT (Ant et al., 2012) have not been translated into policy yet, because the use of genetically modified insects has not been approved in Europe.

Insecticides are primarily used in bait applications, although cover sprays are also applied in some cases. Pyrethroids, spinosyns, neonicotinoids and diamides are registered in Europe and often employed in these applications (EU Pesticides database, 2022). Pyrethroids in particular have been extensively used the last 10-15 years in olive fruit fly control programs in Greece (Minagric, 2022). However, the intense use of insecticides has led to the development of insecticide resistance in *B. oleae*. Moderate to high levels of resistance to pyrethroids have previously been reported in olive fruit fly populations from Greece (Vontas et al., 2011; Margaritopoulos et al., 2008). More recently, in a 9-year long survey of olive fruit fly resistance in Greece, an increase in pyrethroid resistance levels was observed through time, with a clear impact on the efficacy of control applications based on this insecticide, in the field (Kampouraki et al., 2018). In contrast, resistance to other active ingredients, such as spinosad, was found to be low across different populations of Greece.

Resistance to pyrethroid insecticides in *B. oleae* and Tephritidae in more general, has been associated, with detoxification enzymes, while target site resistance mutations on the voltage gated sodium channel have not been identified, as yet (Margaritopoulos et al., 2008; Pavlidi et al., 2018; Vontas et al., 2011). Based on microarray analysis, pyrethroid resistance was associated with the overexpression of two cytochrome P450 genes (Contig00436, probable cytochrome p450 6a23 and Contig02103, probable cytochrome p450 6g2) in *B. oleae* populations from Greece, collected in 2012 and 2014 (Pavlidi et al., 2018). However, their functional link with the phenotype remains elusive and it is not clear if they are capable to metabolise pyrethroids and indeed confer resistance.

Although significant research has been performed on pesticide metabolism and resistance across many insect species, little is known about the exact tissues in which detoxification enzymes, such as CYPs, are localized. The Malpighian tubules (MTs), fat body and midgut are proposed as sites of insecticide detoxification (Beyenbach et al., 2010; Dow and Davies, 2006; Kliot and Ghanim, 2012; Nauen et al., 2022). MTs act as a powerful liver-like detoxification organ and at the same time as a kidney-like secretion system, dedicated to the removal of organic cations and anions from the circulation of the insect (Beyenbach et al., 2010). Previous studies have shown increased expression of cytochrome P450 insecticide metabolizing genes/enzymes in the MTs of resistant insect pests and mosquitoes (Ingham et al., 2014; Wang et al., 2004). In the case of olive fruit flies previous transcriptomic analyses between insecticide-resistant and susceptible olive fly populations have assessed the differential gene expression over the entire organism (Pavlidi et al., 2018, 2013). However, this approach may result in masking effects for some candidate genes, for example if these are differentially expressed specifically in certain tissues, but not at the whole body level.

In this study, we conducted whole body and tissue specific transcriptomics, to identify genes mediating pyrethroid resistance in olive fruit fly field populations from the island of Crete. We also recombinantly expressed four cytochrome P450 genes, that we found over-expressed in resistant populations and characterized their ability to metabolize pyrethroid insecticides. RNAi was used to functionally validate, *in vivo*, the role of one of these P450s in pyrethroid resistance.

## 2. Materials and methods

### 2.1 Insects

Four field populations of *B. oleae* were collected and characterized: Kounoupidiana (Kou17 from Kampouraki et al. 2018), PanagiaA (coordinates 35.115666, 25.338004), PanagiaB (coordinates 35.115108, 25.336146) and Latsida (coordinates 35.266726, 25.592709) from Crete, where olive orchards have been treated with pyrethroid insecticides at a high intensity. In all cases, collected olives infested by *B. oleae*, were transferred into plastic trays and kept at 25 ± 1 °C and 16 h light per day. Heat-sterilised sawdust was used as larval pupation substrate. Pupae were collected daily and transferred to cages. At the day of adult emergence, solid diet (sugar: enzymatic yeast hydrolysate 3:1 w/w) was provided. Water was also provided in a plastic vial, the opening of which was covered with a dental cotton wad. Emerging females were either used for toxicity bioassays or collected and snap frozen in liquid nitrogen for further analysis. Two laboratory strains were also used, as susceptible reference strains: The Lab strain which had not been exposed to insecticides for over 40 years (Tsitsipis and Kontos, 1983), as well as the Hyb, a strain that was created from crosses between filed caught and Lab flies, and has been maintained in the laboratory without exposure to insecticides for over 12 years.

### 2.2 Bioassays

Adult individuals (3-7 days old) were used in toxicity bioassays. Bioassays were carried out by the topical application method. Pyrethroid insecticides alpha-cypermethrin (analytical standard, 45806, Sigma-Aldrich) and deltamethrin (analytical standard, 45423, Sigma-Aldrich) were used. Insecticide doses were prepared by dissolving the insecticide in acetone. Then, 1 ul of the insecticide solution was applied to the dorsal thorax of each adult, using a Hamilton micro syringe of 10 ul volume. Prior to insecticide application, insects were exposed to carbon dioxide for ∼ 12-15 sec to be anaesthetized. At least 20-25 adult individuals were used per dose, and 5-7 doses were applied, including the control (pure acetone). Approximately equal numbers of females and males were used at each dose.

Toxicity tests with the PBO synergist (P450 oxidase inhibitor) were carried out by the same method. More specifically, prior to the application of the insecticide, adult individuals were sprayed with 1 ul of 1000 mg/L PBO solution (PBO 90% technical grade, Sigma-Aldrich, 291102) on the dorsal thorax. This was the highest dose that gave no mortality in susceptible laboratory individuals. The sprayed individuals were allowed to recover for two hours before the insecticide application. Pure acetone was applied instead of insecticide, in the case of controls.

After insecticide application, adult flies were placed in bioassay cages with a constant supply of water and artificial diet and under standard rearing conditions. Mortality was assessed 48 hours after insecticide application. Probit analysis (Finney, 1971) of the results was performed using PoloPC (LeOra Software, Berkeley, CA, USA) to calculate the mean lethal concentration (LC50, lethal concentration-50). The resistance factor (RF, resistance factor) was calculated based on LC50 values as RF = LC50 wild population / LC50 sensitive reference population. The results of the tests involving topical application of insecticides on adult flies are expressed in milligrams of active ingredient per litre of insecticide solution (mg/L).

### 2.3 Transcriptomic analysis

Transcriptome sequencing of whole individuals was performed on the wild strains Kounoupidiana and PanagiaA and the laboratory strain Lab. Four biological replicates from each population, each containing five adult females, 3-5 days old, were sent to the McGill Genome Center (Montreal, Canada). Samples were maintained in RNAlater (Sigma-Aldrich, R0901) until total RNA isolation was performed, using a QIAGEN RNA Extraction Mini Kit (QIAGEN, Germany). The cDNA was synthesized with a Superscript III reverse transcription kit, from 1 ug RNA, and poly dT primers. Illumina TruSeq Library Prep Kit v2 (cat. number: #RS-122-2001) was used to generate libraries. The sequencing reads are available from the Sequence Read Archive (SRA) under the BioProject accession PRJNA891205.

Transcriptome sequencing of malpighian tubules (MT) was performed from the wild flies PanagiaB and the laboratory strain Hyb. Three biological replicates were prepared per each strain and each biological replicate consisted of 7-10 sets of MTs. MTs were dissected out of the adult flies on ice, in PBS 1X and collected in DNAse free Eppendorf tubes, snap frozen in liquid nitrogen and stored at -80°C until RNA extraction. Total RNA was extracted using the TRI Reagent (Molecular Research Center, Inc., TR 118) according to the manufacturer’s instructions and RNA pellets were re-suspended in RNase-free DEPC water. Afterwards, the samples were DNase-treated using DNAse I (Ambion DNase I (RNase-free), AM2222) in order to remove any contaminating DNA. RNA concentration and purity (OD, A260/A280 and A260/A230) were measured in a Nanodrop ND-1000 Spectrophotometer (NanoDrop Technologies, Wilmington, DE, USA) using 1 ul of RNA. The ratio between the absorbance at 260 nm and 280 nm was used to evaluate purity; we assumed ratios between 1.9 and 2.0 to be pure enough for the RNAseq analysis. RNA integrity was checked by electrophoresis of 500 ng extracted RNA on a 1.5% agarose gel.

Approximately 1.5ug of each purified RNA sample, from each one of the three biological replicates for Hyb and PanagiaB strains were sent to Macrogen, Inc. (Korea) for mRNA paired end library construction with the Illumina Truseq stranded mRNA sample preparation kit, following the manufacturer’s instructions. Each library was sequenced on the Illumina platform with the paired-end method and a read length of 100 bp. The sequencing reads are available from the Sequence Read Archive (SRA) under the BioProject accession PRJNA891205.

Reads were first quality-trimmed using trimmomatic (Bolger et al., 2014), in order to remove sequencing adapters and low-quality bases. Trimmed reads were then mapped on the publicly available *B. oleae* reference genome ((Bayega, 2021), GCF_001188975.3) using the Hisat2 short read aligner (Kim et al., 2019), and read counts for each of the predicted genes in the official gene set were calculated with Feature Counts (Liao et al., 2014) at the gene level. EdgeR (Robinson et al., 2010) was used to find genes that were significantly (FDR <1e-03) differentially expressed with a fold change > 4 (log_2_FC > 2).

Determining whether *B. oleae* genes were full-length was done using BLAST. More specifically, each *B. oleae* predicted protein was searched against the Uniref50 protein database with an e-value cut-off of 1e-25. Self-hits were first excluded and by using custom Perl scripts we searched for hits covering >90% of the *B. oleae* protein (query), as well as the database protein (subject). This approach ensured that the *B. oleae* protein had an end-to-end match with a protein from another species.

### 2.4 Quantitative PCR transcript analysis

Transcriptomic data of four CYP genes were validated using quantitative reverse transcription PCR (RT-qPCR).

Total RNA from the whole bodies (four insects in each biological replicate/sample) and from MT (7-10 MTs in each sample/biological replicate) was isolated with TRIzol (Invitrogen, 15596026). Isolated RNA samples were treated with DNase I (Thermo Scientific, EN0521). Reverse transcription was performed on 0.5 or 1 ug of isolated RNA for MTs and whole bodies respectively, using Reverse Transcriptase (MINOTECH RT, 801-1) and Oligo dT (20) primers. PCR reactions of 25 ul were performed on a MiniOpticon two-colour Real-Time PCR detection system (BioRad), using 0.20 uM of primers and KapaSYBR FAST qPCR master mix (Kapa Biosystems).

Primers for PCR were designed using the Primer-BLAST online analysis software (http://www.ncbi.nlm.nih.gov/tools/primer-blast/). These primers are listed in Table S1. Experiments were performed in at least 3 biological replicates for each gene. The expression level of each target gene was normalized against the 40S ribosomal protein (GAKB01005984.1) and beta-actin (GAKB01001968.1) reference genes (Pavlidi et al., 2018). The results were analysed using the relative quantification approach through the Comparative Ct method (Pfaffl, 2001).

### 2.5. Cloning, co-expression of P450s with *Md*CPR and preparation of membranes

The sequences encoding the target cytochrome CYP genes were optimized for codon usage in bacteria and synthesized in AZENTA in PUC_GW vector. The primers used for the subcloning of these genes into the pCW-OmpA2 vector (about 40 copies per cell) are listed in Table S1. *Md*CPR sequence was optimized for codon usage in bacteria and synthesized in TWIST in pCDFDuet-1 (20-40 copies per cell).

For protein expression, the previously described strategy (Riga et al., 2014) was followed using *E. coli* BL21STAR cells co-transformed with the pCW-OmpA2-P450 and the pCDFDuet-1-MdCPR. Transformed cells were grown in terrific broth medium with ampicillin and streptomycin selection until the optical density at 600 nm reached ∼ 0.8-0.9 cm^-1^, whereupon the heme precursor δ-aminolaevulinic acid (ALA) was added to a final concentration of 1 mM. Induction was initiated with the addition of isopropyl-1-thio-β-D glucopyranoside (IPTG) to a final concentration of 0.5 mM. Spheroplasts are prepared by adding 1x TSE buffer (0.1 M Tris acetate, pH 7.6, 0.5 M sucrose, 0.5 mM EDTA) containing 0.25 mg/ml lysozyme to the cell pellet and gentle mixing for 60 min at 4°C. The solution is centrifuged at 2800 g for 25 min at 4°C and the spheroplast pellet is resuspended in spheroplast resuspension buffer (0.1 M potassium phosphate buffer, pH 7.6, 6 mM magnesium acetate, 20% glycerol) containing 0.1 mM dithiothreitol (DTT), 1 mM phenylmethanesulfonylfluoride (PMSF), 1 mg/ml aprotinin and 1 mg/ml leupeptin. Then, the suspension is sonicated and centrifuged at 13000 rpm for 25 min at 4° C. The supernatant was collected and ultracentrifuged at 180,000 g for 1 h, at 4°C. Membrane preparations were diluted in 1x TSE buffer and stored at -80 °C. Membrane aliquots are assayed for total protein concentration (Bradford assay with BSA standards), CO-spectrum (Omura and Sato, 1964), and CPR activity by monitoring cytochrome c reduction (Strobel and Dignam, 1978).

### 2.6 Analysis of insecticide metabolism

We examined evidence for metabolism of three pyrethroid insecticides, deltamethrin, alpha-cypermethrin and lambda-cyhalothrin, and two neonicotinoid insecticides, thiacloprid and acetamiprid (Sigma Aldrich). Stock concentrations of pyrethroid and neonicotinoid insecticides were prepared and diluted in acetonitrile and methanol, respectively. Standard reactions contained a final organic solvent concentration of 2,5 % (v/v) with 25 uM of each insecticide, 50 pmol of each P450 and 500 pmol Agcytb5 in 100 ul Tris-HCl buffer (0.2 M, pH 7.4), containing 0.25 mM MgCl_2_. The incubation was performed in the presence and absence of an NADPH generating system: 1 mM glucose-6-phosphate (Sigma Aldrich) 0.1 mM NADP+ (Sigma Aldrich) and 1 unit/ml glucose-6-phosphate dehydrogenase (G6PDH-Sigma Aldrich). Reactions were incubated at 30 °C, 1250 rpm oscillation and stopped at 0- and 2-hour time points using 100 ul acetonitrile and stirred additionally for 30 min. Finally, the quenched reactions were centrifuged at 10,000 rpm for 10 min and the supernatant was transferred to HPLC vials, with 100 ul of the supernatant loaded for HPLC analysis. Reactions were performed in triplicate and compared against a negative control with no NADPH regenerating system to calculate substrate depletion. Insecticides were separated on a UniverSil HS C18 (250 mm 5 um) reverse phase analytical column (Fortis) and a mobile phase optimized for each insecticide (Table S2). Insecticides were quantified by peak integration (Chromeleon, Dionex). The absorbance and the elution time of each insecticide are also summarized in Table S2.

### 2.7 dsRNA synthesis and RNAi of P450 genes

For double-stranded RNA (dsRNA) synthesis, PCR products of the target CYP gene and LACZ as control, were generated using primers with T7 promoter sequence (Table S1). The newly synthesized PCR products were then purified with NucleoSpin Gel and PCR Clean-up (Macherey–Nagel), which were subsequently used as template for the synthesis of dsRNA molecules. Synthesis was performed with the HiScribe T7 High Yield RNA Synthesis Kit (New England Biolabs), according to the manufacturer’s instructions. After synthesis, the produced dsRNA was purified by phenol: chloroform extraction and recovered by ethanol precipitation. The quality of the products was checked on 1,5 % w/v agarose gel and the concentration of each sample was determined using a small volume spectrophotometer.

The dsRNA was injected into 2-4 days old adults according to (Huang et al., 2015) methodology. Briefly, after the insects were left on ice until anesthetized, approximately 1 ug of dsRNA was injected into each individual, between the first and second abdominal segments. Injections were performed using the Nanoject II Auto-Nanoliter Injector (Drummond Scientific, Broomall, PA, USA). Approximately 30 individuals were injected in each replicate. Mortality was recorded, 24h after the injections, and it was in all cases <10%. Injected individuals were maintained under conditions of constant temperature, photoperiod and food supply. Target-gene expression levels were evaluated in 5 individuals collected 24 h after dsRNA insertion, by quantitative RT-PCR (as described above). The remaining individuals, about 25 in each replicate, were used in diagnostic toxicity bioassays (as described above), with the insecticide deltamethrin, in a concentration corresponding to the LC10 of the population used.

## 3. Results

### 3.1 Resistance levels of *Bactrocera oleae* field populations from Crete

Olive flies were collected at four different locations (Kounoupidiana (toxicity data from (Kampouraki et al., 2018)), PanagiaA, PanagiaB and Latsida) in Crete and their resistance to alpha-cypermethrin and deltamethrin assessed by topical application of the insecticides. According to the established classification of (Torres-Vila et al., 2002a, 2002b) the Kounoupidiana, PanagiaA and PanagiaB populations showed very high resistance to alpha-cypermethrin, compared to the susceptible Lab strain, with resistance ratios ranging from 222-fold in PanagiaB to 331-fold in Kounoupidiana (Table 1). Deltamethrin toxicity was also tested in PanagiaB and Latsida populations, and was found more toxic than alpha-cypermethrin. Resistance ratios, compared to the susceptible Hyb strain, were 10-fold for Panagia B and 23-fold for Latsida.

**Table 1.**
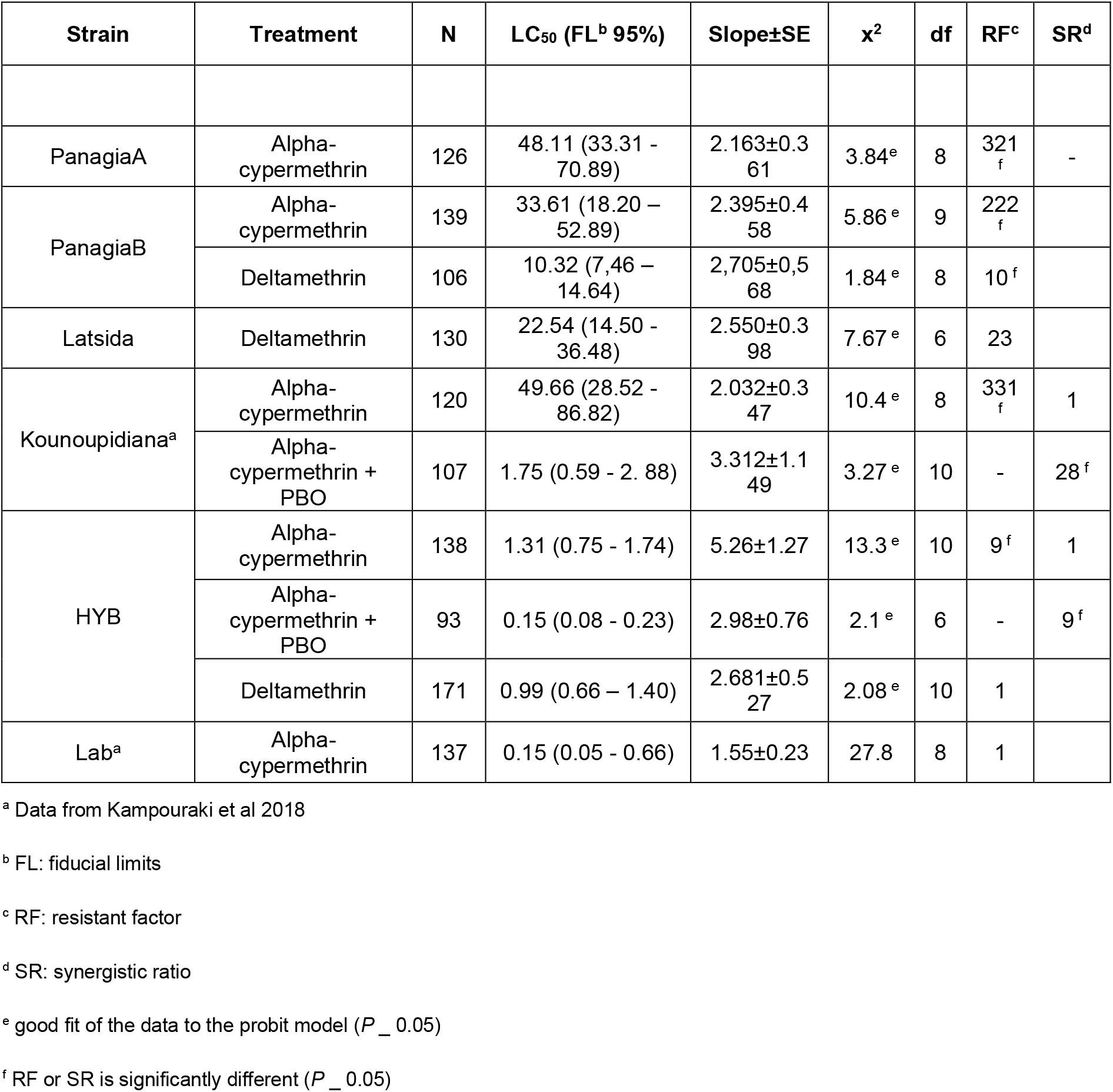
Susceptibility of *B. oleae* field and laboratory (Lab and Hyb) populations to alpha-cypermethrin and deltamethrin insecticides using topical application bioassays, and synergistic effect of PBO on the toxicity of alpha-cypermethrin to the field population Kounoupidiana and the laboratory population HYB.

| Strain | Treatment | N | LC<sub>50</sub> (FL<sup>b</sup> 95%) | Slope±SE | χ<sup>2</sup> | df | RF<sup>c</sup> | SR<sup>d</sup> |
| --- | --- | --- | --- | --- | --- | --- | --- | --- |
| PanagiaA | Alpha-cypermethrin | 126 | 48.11 (33.31 - 70.89) | 2.163±0.361 | 3.84<sup>e</sup> | 8 | 321<sub>f</sub> | - |
| PanagiaB | Alpha-cypermethrin | 139 | 33.61 (18.20 – 52.89) | 2.395±0.458 | 5.86<sup>e</sup> | 9 | 222<sub>f</sub> |  |
|  | Deltamethrin | 106 | 10.32 (7.46 – 14.64) | 2,705±0,568 | 1.84<sup>e</sup> | 8 | 10<sup>f</sup> |  |
| Latsida | Deltamethrin | 130 | 22.54 (14.50 - 36.48) | 2.550±0.398 | 7.67<sup>e</sup> | 6 | 23 |  |
| Kounoupidiana<sup>a</sup> | Alpha-cypermethrin | 120 | 49.66 (28.52 - 86.82) | 2.032±0.347 | 10.4<sup>e</sup> | 8 | 331<sub>f</sub> | 1 |
|  | Alpha-cypermethrin + PBO | 107 | 1.75 (0.59 - 2.88) | 3.312±1.149 | 3.27<sup>e</sup> | 10 | - | 28<sup>f</sup> |
| HYB | Alpha-cypermethrin | 138 | 1.31 (0.75 - 1.74) | 5.26±1.27 | 13.3<sup>e</sup> | 10 | 9<sup>f</sup> | 1 |
|  | Alpha-cypermethrin + PBO | 93 | 0.15 (0.08 - 0.23) | 2.98±0.76 | 2.1<sup>e</sup> | 6 | - | 9<sup>f</sup> |
|  | Deltamethrin | 171 | 0.99 (0.66 – 1.40) | 2.681±0.527 | 2.08<sup>e</sup> | 10 | 1 |  |
| Lab<sup>a</sup> | Alpha-cypermethrin | 137 | 0.15 (0.05 - 0.66) | 1.55±0.23 | 27.8 | 8 | 1 |  |

<sup>a</sup> Data from Kampouraki et al 2018
<sup>b</sup> FL: fiducial limits
<sup>c</sup> RF: resistant factor
<sup>d</sup> SR: synergistic ratio
<sup>e</sup> good fit of the data to the probit model ( $P \geq 0.05$ )
<sup>f</sup> RF or SR is significantly different ( $P \geq 0.05$ )

### 3.2 Toxicity assays with the PBO synergist

Toxicity bioassays with the PBO synergist, which acts as a P450 inhibitor, were carried out in the Kounoupidiana population, which was the most resistant to pyrethroids (alpha-cypermethrin), and in the laboratory strain Hyb. These bioassays were conducted in order to determine whether the observed resistance was, at least in part, due to detoxification by P450s. In both populations, the PBO synergist enhanced the effect of alpha-cypermethrin.

More specifically, in the Kounoupidiana population the sensitivity to the insecticide increased by 28-fold, indicating a high synergistic effect, while in the Hyb population the increase in sensitivity was much lower (by 8-fold) (Table 1). In addition, after PBO application in the Kounoupidiana population, the mean LC_50_ value dropped from 49.66 mg/L to 1.74 mg/L, which is very close to the LC_50_ of the Hyb population (1.31 mg/L) without PBO application. This drop in resistance strongly suggests that P450-related detoxification is involved in resistance of this population to pyrethroids (alpha-cypermethrin).

### 3.3 Whole-body comparative transcriptomic analysis in pyrethroid resistant and susceptible *B. oleae* populations

Comparison of whole-body gene expression profiles was performed between the highly resistant Kounoupidiana and PanagiaA populations against the susceptible Lab strain. Four replicates were sequenced for each strain, yielding a total of 1,643 million reads (Table S3). Principal components analysis (PCA) showed that replicates of these populations were separated from each other (Figure S1), thus ensuring the validity of the subsequent comparative analyses.

Differential expression analysis identified 150 genes significantly (|log_2_FC| > 2 and FDR < 0.05) up-regulated in at least one of the two resistant populations (Kounoupidiana/PanagiaA), compared to the susceptible Lab strain (Table S4). More specifically, we found 138 significantly up-regulated genes in Kounoupidiana and 53 in PanagiaA, 41 of which are commonly up-regulated in the two populations.

Among the over-expressed genes, we found 11 cytochrome P450s and 4 UDP-glycosyltransferases (UGTs) (Figure 1A, Table S5). Of these, five P450s showed over-expression in both Kounoupidiana and PanagiaA populations. These P450s are: *Cyp6A61* (similar to *D. melanogaster Cyp6A23*, over-expressed by 43.3-fold in Kounoupidiana and 24.9-fold in PanagiaA), *Cyp3162A1* (similar to *D. melanogaster Cyp313A4*, over-expressed by 20.6-fold in Kounoupidiana and 17.6-fold in PanagiaA), *Cyp6G6* (similar to *D. melanogaster Cyp6G2*, over-expressed by 12.9-fold in Kounoupidiana and 9.2-fold in PanagiaA) and its paralog *Cyp6G27* (over-expressed by 11.1-fold in Kounoupidiana and 8-fold in PanagiaA), and *Cyp9B7* (similar to *D. melanogaster Cyp9B2*, 7-fold in Kounoupidiana and 6.7-fold in PanagiaA). UGTs, on the other hand, were up-regulated only in Kounoupidiana and not in PanagiaA (Figure 1A).

**Figure 1.**
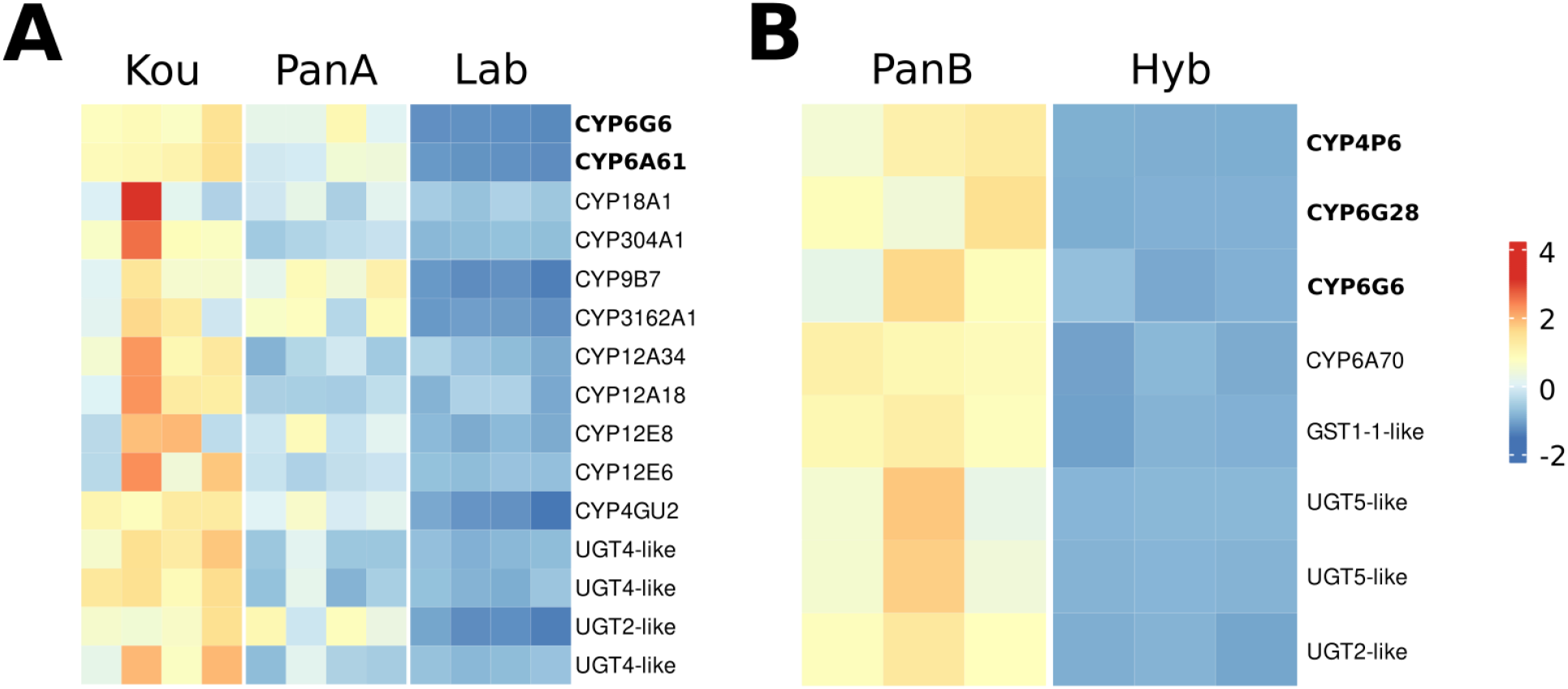
Expression levels of all significantly over-expressed detoxification genes in the two transcriptomics data sets. **(A)** Expression levels in whole body samples from the resistant Kounoupidiana (Kou) and PanagiaA (PanA) populations against the susceptible Lab population. (**B)** Expression levels in malpighian tubules from the resistant PanagiaB (PanB) population against the susceptible Hybrid (Hyb) population. Expression values are row-normalized z-scores. In bold are the P450s that were selected for *in vitro* metabolism assays.

The genomic locus containing *Cyp6G6* seems to have three copies of this gene (*Cyp6G6, Cyp6G24*, and *Cyp6G27*). *Cyp6G6* and *Cyp6G27* appear to be significantly up-regulated in the pyrethroid-resistant populations (based on our RNAseq results-Table S5). These two CYPs are 91% similar to each other on the amino acid level, while *Cyp6G24* is much more divergent (∼55% similar to either 6G6/6G27). However, upon closer examination, only *Cyp6G6* appears to be uniformly transcribed and could thus be implicated in resistance to alpha-cypermethrin.

The over-expression of *Cyp6G6* (LOC106618206-00002) and *Cyp6A61* (LOC106620242), both of which belong to the CYP6 family, was further validated by qRT-PCR (Figure 2A). *Cyp6A61* was found over-expressed (compared to the Lab strain) by 12.5 and 6.9-fold in the resistant Kounoupidiana and PanagiaA populations, respectively. Likewise, *Cyp6G6* was found over-expressed by 5.5 and 8.5-fold in Kounoupidiana and PanagiaA, respectively (Figure 2A)

**Figure 2.**
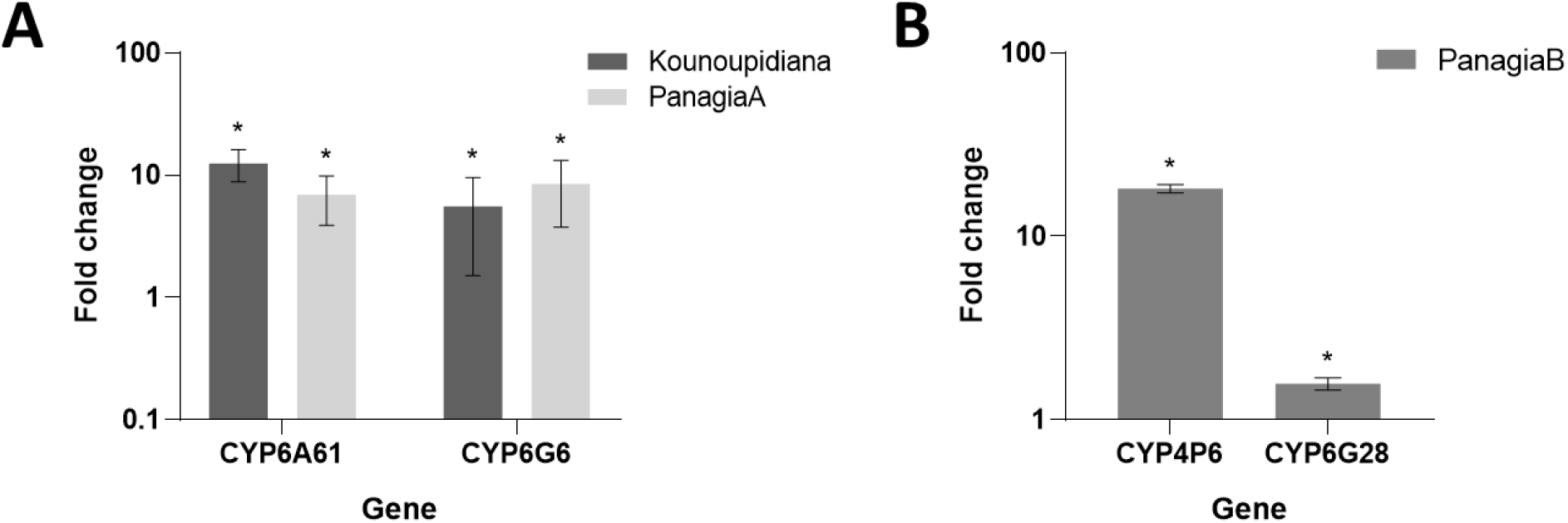
qPCR analysis of selected CYP transcripts identified as differentially expressed in RNA-Seq **(A)** *Cyp6A61* and *Cyp6G6* genes in whole bodies of Kounoupidiana and PanagiaA field populations relative to Lab laboratory strain, and **(B)** CYP4P6 and *CYP6G28* genes in MTs of PanagiaB field population relative to Hyb laboratory strain. Error bars represent the SD of the mean of at least three independent biological replicates. An Asterisk (*) indicates a statistically significant difference (P<0.01) of relative expression levels.

### 3.4 Comparative transcriptomic analysis in malpighian tubules from pyrethroid resistant and susceptible *B. oleae* populations

Differential gene expression was also studied in the malpighian tubules of the resistant PanagiaB population and the susceptible Hybrid (Hyb) population, which has been maintained under laboratory conditions for at least 150 generations. A total of 378 million short reads were generated from PanagiaB and Hyb (Table S6). A principal components analysis (PCA) using the expression levels of all expressed genes showed that the replicates of the two populations clustered separately from each other (Figure S2), indicating that their pairwise comparison is valid. Several detoxification genes were found up-regulated in PanagiaB. More specifically, among the 123 genes that were up-regulated (Table S7), 19 of which were also over-expressed in the whole-body analysis, we identified four P450s from the CYP3 and CYP4 clans (Figure 1B, Table S8).

*Cyp4P6* (similar to *D. melanogaster Cyp4P1*) is the most up-regulated P450 (93.7-fold in PanagiaB) in this tissue-specific dataset (Table S8). This P450 is highly transcribed in the resistant strain, while it is virtually absent from the susceptible one. Furthermore, *Cyp6G28*, similar to the *D. melanogaster Cyp6G1*, is also significantly up-regulated in the resistant strain (15.9-fold).

The remaining three up-regulated P450s include: the *Cyp6G6* (10.1-fold) and *Cyp6G27* (15-fold) (both are similar to *Cyp6G2* from *D. melanogaster*), as well as the *Cyp6A70* (4.4-fold) (similar to *Cyp6A13* from *D. melanogaster*). *Cyp6G6* has been previously associated to resistance against pyrethroid insecticides (contig02103 in (Pavlidi et al., 2018))

Quantitative PCR (qRT-PCR) was used to validate the up-regulation of two of the identified P450s, in the malpighian tubules of the resistant strain. CYP4P6 was found over-expressed by 18.20-fold, while *Cyp6G28* was lowly, but significantly over-expressed at 1.57-fold (Figure 2B).

Among the over-expressed genes, we also identified one GST: locus name (LOC106615161), which is mostly similar to GST 1-1 from various Tephritid species. It has a length of 215 amino acids and appears to be full-length. Moreover, its transcription is uniform and follows the gene structure. As for the three UGTs, two of them (LOC118679823 and LOC106623379) appear to be full-length and both are UGT5-like. The third UGT (LOC118682404) is fragmented, apparently having a N-term truncation. The two full-length UGTs, are highly overexpressed in the resistant strain, at almost 210-fold for LOC118679823 and 32-fold for LOC106623379 (Table S8).

### 3.5 Heterologous expression of *B. oleae* P450s in *E. coli*

The *Cyp6A61, Cyp6G6, Cyp4P6* and *Cyp6G28* genes, which showed increased transcription levels in resistant populations, were chosen for expression in *E. coli. Cyp6A61* was the most over-expressed detoxification gene, in the whole-body transcriptomic analysis; *Cyp6G6* was found over-expressed in the analyses of both whole bodies and MTs; and Cyp*4P6* and *Cyp6G28* were the two detoxification genes most highly overexpressed in MTs of the resistant population. All four P450swere co-expressed with the *Musca domestica* CPR in *E. coli* membranes, producing catalytically active monooxygenase complexes (Omura and Sato, 1964). *An. gambiae* cytb5 was also used to enhance the P450 catalytic activity. All membrane preps produced contained high levels of P450s (up to 175 nmol/L) (Table S9) and showed the characteristic CO-reduced spectrum, which is indicative of an active P450 (Figure S3). In all cases, CPR activity against cytochrome c was confirmed, as shown in Table S9.

### 3.6 Insecticide metabolism

The metabolic activity of these P450s towards different pyrethroid insecticides (deltamethrin, alpha-cypermethrin and lambda-cyhalothrin Type II (cyano)) and neonicotinoids (thiacloprid and acetamiprid) was tested by measuring insecticide depletion in the presence and absence of NADPH (Table 2). For the three pyrethroids that were tested, only deltamethrin and alpha-cypermethrin were metabolized, by at least one P450. More specifically, CYP6G6 was able to metabolize 33,4% of the parental deltamethrin, after 2h incubation, compared to CYP6G28 and CYP4P6, both of which metabolized at lower levels the parental deltamethrin (∼24% and 17%, respectively). CYP6A61 was the only P450 from the panel which showed activity against alpha-cypermethrin. CYP6A61 was able to metabolize 13.2% of the parental alpha-cypermethrin. None of the P450s showed activity against neonicotinoid insecticides.

**Table 2.** Pyrethroid and neonicotinoid metabolism by *B. oleae* P450s

| P450s | % Insecticide depletion |  |  |  |  |
| --- | --- | --- | --- | --- | --- |
|  | Pyrethroids type II |  |  | Neonicotinoids |  |
|  | Deltamethrin | lambda-cyhalothrin | alpha-cypermethrin | Thiacloprid | Acetamiprid |
| CYP6G28 | 23,75 ± 0,04 | nd<sup>a</sup> | nd<sup>a</sup> | nd<sup>a</sup> | nd<sup>a</sup> |
| CYP4P6 | 17 ± 0,15 | nd<sup>a</sup> | nd<sup>a</sup> | nd<sup>a</sup> | nd<sup>a</sup> |
| CYP6G6 | 33,4 ± 1,8 | nd<sup>a</sup> | nd<sup>a</sup> | nd<sup>a</sup> | nd<sup>a</sup> |
| CYP6A61 | nd<sup>a</sup> | nd<sup>a</sup> | 13,2 ± 0,68 | nd<sup>a</sup> | nd<sup>a</sup> |

<sup>a</sup> Non detected

### 3.7 Knockdown of CYP6A61 by RNAi

Given that *Cyp6G6* was found significantly up-regulated in both transcriptomic analyses of whole bodies and MTs, and able to metabolize deltamethrin *in vitro*, we used RNAi to validate its role in resistance *in vivo*. The field resistant population Latsida was used for the RNAi experiments. The expression levels of *Cyp6G6* were evaluated in dsCyp6G6-injected adults, collected 24 h after injections. As shown in Figure 3A, *Cyp6G6* expression levels were reduced in dsCYP6G6 injected individuals by ∼65%, compared to dsLACZ (dsRNA for the non-endogenous *LacZ*-) injected individuals (controls).

**Figure 3.**
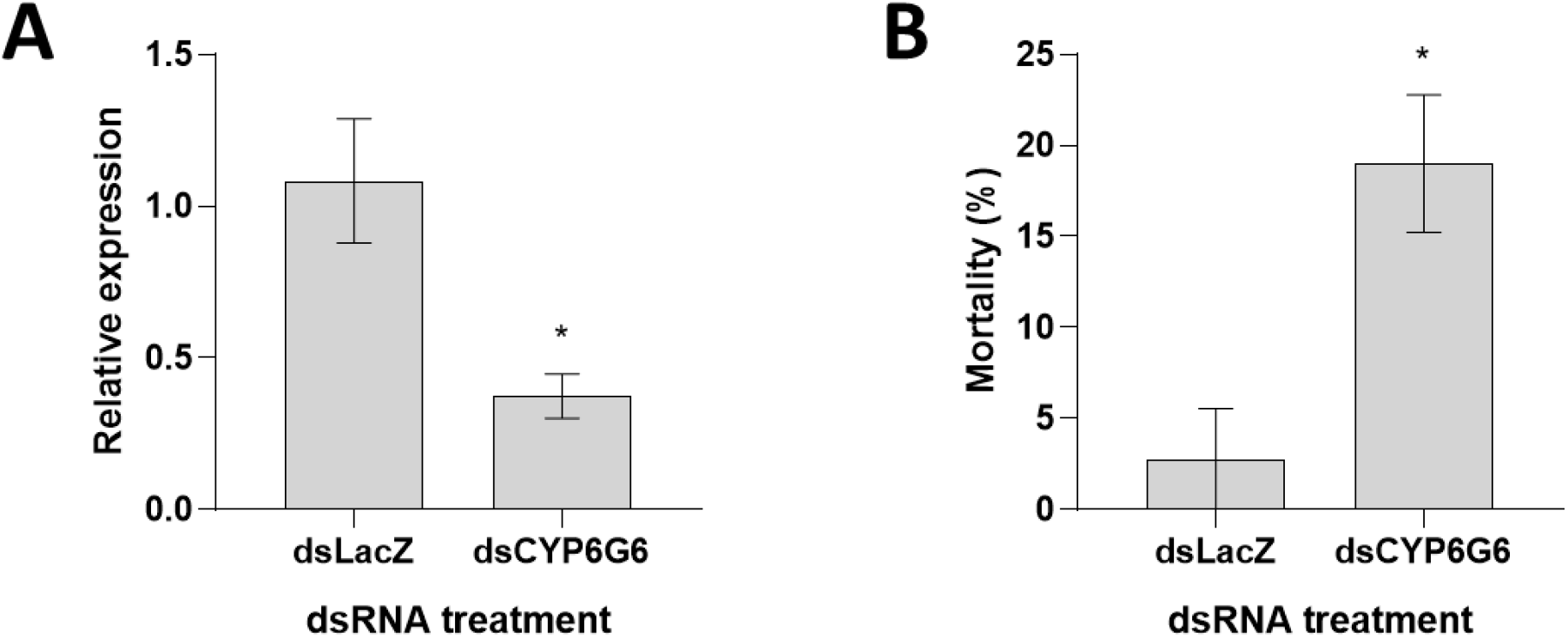
Efficiency of *Cyp6G6* silencing in the Latsida population and its effect on resistance to deltamethrin. **(A)** Expression levels of *Cyp6G6* in individuals injected with dsRNA targeting *LacZ* (as control) or *Cyp6G6*, 24 hours post injection. **(B)** Mortality (%) of dsLacZ-injected and dsCyp6G6-injected *B. oleae* 48 hours after alpha-cypermethrin treatment. Error bars represent the SE of the mean of three independent biological replicates. An Asterisk (*) indicates a statistically significant difference in expression levels (A) or percentage of mortality (B) between treatments (P<0.05).

Diagnostic toxicity bioassays with deltamethrin in injected adults, were performed 72 hours post injections. Reduction in the expression of *Cyp6G6* significantly increased sensitivity to deltamethrin raising the mean mortality to 19% compared to the 2.8% mortality observed in the control dsLACZ individuals (Figure 3B).

## 4. Discussion

Olive plantations on the island of Crete, a major oil-producing region of Greece, have been intensely treated with pyrethroids for over 10 years to prevent destruction of olive oil production from *B. oleae*. In this study, field and laboratory susceptible *B. oleae* populations were subjected to toxicity bioassays with alpha-cypermethrin and deltamethrin, and the results confirmed the presence of highly pyrethroid resistant populations in Crete. These results are in line with previous reports of the presence of highly pyrethroid resistant populations in the same region (Kampouraki et al., 2018), and in fact reveal that resistance has increased over the years.

Despite the implementation of evidence-based rotation strategies in Crete and the extensive and almost exclusive use of spinosad by the national control program in the last period (2019-2021), pyrethroid resistance remains high, possibly due to the high number of cover pyrethroid sprays by individual farmers for therapeutic systemic applications.

Resistance to alpha-cypermethrin was substantially suppressed by PBO, in toxicity assays, indicating that P450s play an essential role in pyrethroid resistance, which is in agreement with previous findings (Pavlidi et al., 2018).

In order to identify the mechanisms of pyrethroid resistance in *B. oleae*, we undertook a comparative transcriptomic analysis, both of whole bodies and malpighian tubules, a tissue considered a main site of insecticide metabolism, between pyrethroid resistant field populations from Crete and susceptible laboratory strains. Each of these analyses revealed a number of overexpressed detoxification genes, primarily CYPs. A small number of UGTs and GSTs were also found overexpressed.

From the P450s identified as over-expressed in the whole-body analysis, *Cyp6A61, Cyp6G6, Cyp3162A1* and *Cyp9B7* were also reported over-expressed in pyrethroid resistant field populations from the same region (Crete) in 2018 (Pavlidi et al., 2018), indicating their consistent over-expression. The level of over-expression of these genes has also increased since 2018 (Pavlidi et al., 2018), which is in line with the observed increase in pyrethroid resistance levels. Furthermore, *Cyp6G6* and *Cyp9B7* were previously found up-regulated in a malathion resistant strain of *Bactrocera dorsalis*, while *Cyp6A61* was found down-regulated in the same strain (Jing et al., 2020).

The Malpighian tubule transcriptomic analysis showed 123 genes over-expressed in the pyrethroid resistant population, a 15% of which were also over-expressed in the whole-body analysis. Importantly though the Malpighian tubule analysis revealed a largely different subset of detoxification genes to be up-regulated compared to the whole-body analysis, with only *Cyp6G6* being commonly up-regulated. This could be related to the different populations/strains used in each comparison (Kounoupidiana and PanagiaA vs Lab for whole-body and PanagiaB vs Hyb for MTs), but it could also show a tissue specific expression or differential expression, that is masked when using whole-bodies.

We provided further support for the role of four of the over-expressed P450s in pyrethroid resistance, by heterologously expressing them and showing their ability to metabolise deltamethrin and/ or cypermethrin *in vitro*. We also showed that down-regulation of *Cyp6G6* through RNAi silencing results in a small, but significant reduction in resistance *in vivo*. The relatively small effect observed could is related to the complex genetic basis of resistance involving the over-expression of multiple detoxification enzymes and their functional redundancy in the resistance phenotype.

Our herein developed in vitro assays support the role of Cyp6G28, Cyp4P6, Cyp6G6 and Cyp6A61 in pyrethroid detoxification. At the same time the *in vitro* assays suggest that these enzymes are not involved in the detoxification of other classes of insecticides, such as the neonicotinoids thiacloprid and acetamiprid, also used for *B. oleae* control, as all of the enzymes were found inactive against these molecules. This indicates that these enzymes may have a specialized role in pyrethroid resistance, similar to CYP6A51 from *Ceratitis capitata*, which was found to metabolize only deltamethrin and lambda-cyhalothrin, but not spinosad and malathion (Tsakireli et al., 2019) (Tsakireli et al., 2019), as well as the CYP392A11 and CYP392E10 from *Tetranychus urticae* which metabolize only METIs and spirodiclofen, respectively (Demaeght et al., 2013; Riga et al., 2020, 2015, 2014) (Demaeght et al., 2013; Riga et al., 2020, 2015, 2014). This is in contrast to other P450s, which have a broader catalytic role, such as the CYP6G1 from *Drosophila melanogaster*, which is capable of metabolizing both the organochlorine DDT and the neonicotinoid imidacloprid (Joussen et al., 2008) (Joussen et al., 2008), or the CYP6CM1 from *Bemisia tabaci*, which is capable of metabolizing neonicotinoids, pyridine azomethin and pyriproxyfen (Karunker et al., 2009; Nauen et al., 2013; Roditakis et al., 2011) (Karunker et al., 2009; Nauen et al., 2013; Roditakis et al., 2011).

In conclusion, this study has established that four olive fruit fly cytochrome P450s are overexpressed in pyrethroid resistant field populations and they are capable to metabolize pyrethroids and confer resistance. This information is valuable towards the development of molecular diagnostics and evidence based insecticide resistance management strategies.

## Acknowledgements

We thank Dr. Mark Paine (LSTM) for kindly providing the expression vector PCWompA, Aggeliki Karataraki (Directorate of Rural Economy and Veterinary, Regional unit of Lassithi, Crete) and Eleni Malandraki (Directorate of Rural Economy and Veterinary, Regional unit of Chania, Crete) for providing *B*.*oleae* field populations (Latsida and Kounoupidiana respectively). P.I. and V.K. were supported by the Operational Programme “Human Resources Development, Education and Lifelong Learning 2014-2020” (co-financed by Greece and the European Social Fund) in the context of the project “Using bioinformatics and biotechnology, in order to study insecticide resistance in the olive fruit fly, *Bactrocera oleae*” (MIS 5052108). A.K. was supported by the General Secretariat for Research and Technology (GSRT) and the Hellenic Foundation for Research and Innovation (HFRI) in the context of the action “1st Proclamation of Scholarships from ELIDEK for PhD Candidates” (Scholarship Code: 2283)

## Supporting Information

### Tables

**Table S1.**
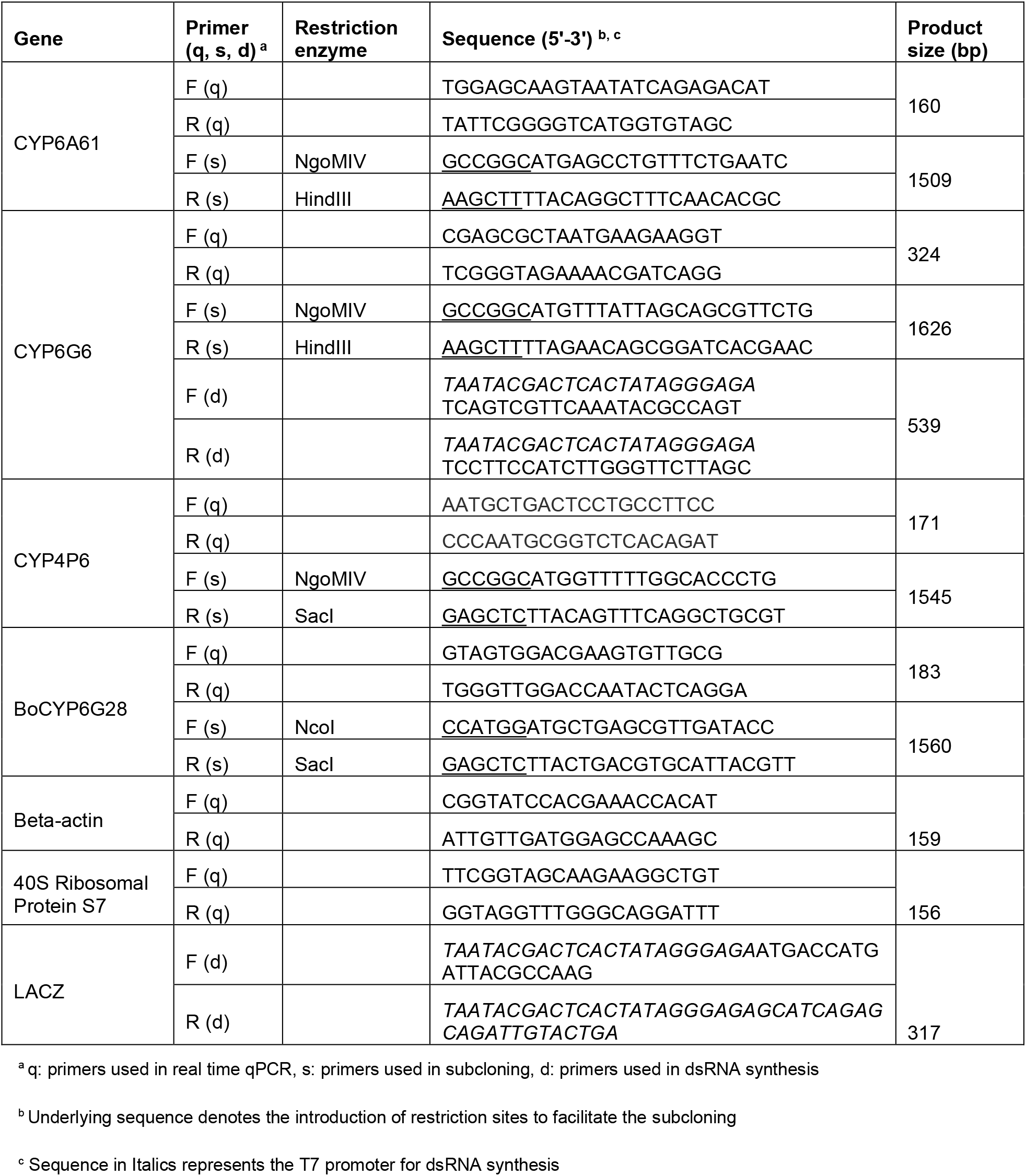
Primers used in this study

**Table S2.**
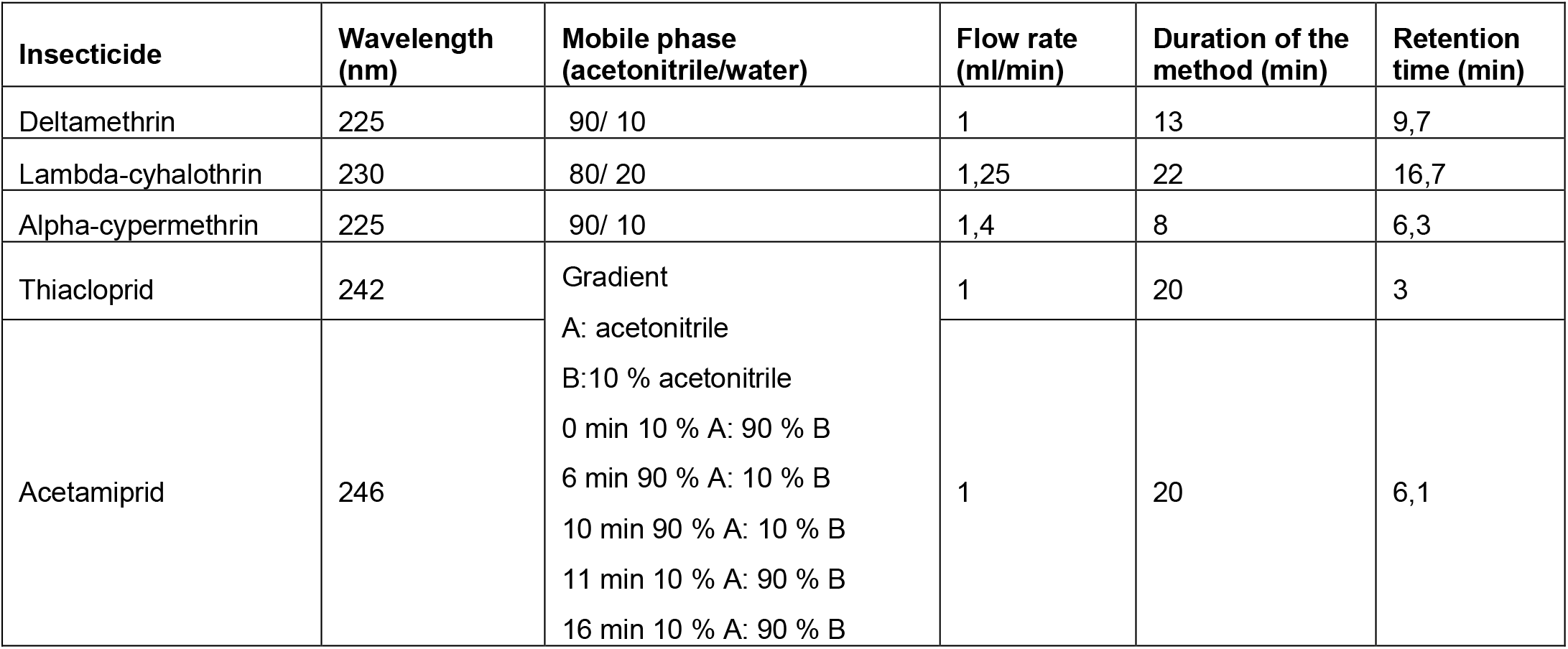
Analytical HPLC conditions for insecticide peaks detection

**Table S3.**
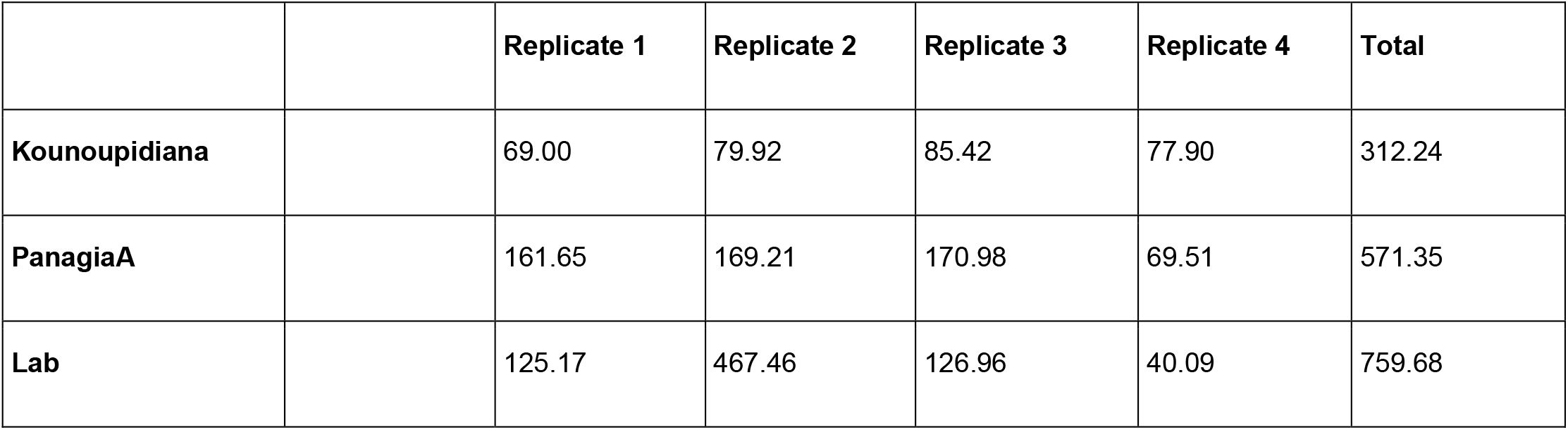
Number of RNAseq reads (million reads) generated for each sample, for the whole-body study.

**Table S5.** Significantly (FDR <0.001 and log_2_FC >2) up-regulated detoxification genes in the four whole-body samples.

| Gene id | Up-regulation (log<sub>2</sub>FC) |  |  |  | Annotation |
| --- | --- | --- | --- | --- | --- |
|  | Kou-Lab<sup>1</sup> | PanA-Lab<sup>1</sup> |  |  |  |
| LOC106618206-00002 | 3,693 | 3,198 |  |  | CYP6G6<sup>2</sup> |
| LOC106618206-00001 | 3,469 | 3,005 |  |  | CYP6G24<sup>3</sup> |
| LOC106615810 | 2,132 | N/A |  |  | CYP18A1 |
| LOC118680302 | 2,219 | N/A |  |  | CYP4GU2 |
| LOC106618459 | 2,808 | 2,745 |  |  | CYP9B7 |
| LOC106622341 | 2,035 | N/A |  |  | CYP12A34 |
| LOC106622342 | 2,291 | N/A |  |  | CYP12A18 |
| LOC106622736 | 2,395 | N/A |  |  | CYP12E6 |
| LOC106622732 | 2,37 | N/A |  |  | CYP12E8 |
| LOC106617790 | 3,361 | N/A |  |  | CYP304A1 |
| LOC106622034 | 4,365 | 4,138 |  |  | CYP3162A1 |
| LOC106620242 | 5,436 | 4,638 |  |  | CYP6A61<sup>4</sup> |
| LOC118682404 | 2,204 | N/A |  |  | UGT2-LIKE |
| LOC106623376 | 3,253 | N/A |  |  | UGT4-LIKE |
| LOC118682405 | 3,114 | N/A |  |  | UGT4-LIKE |
| LOC118680791 | 2,737 | N/A |  |  | UGT4-LIKE |

<sup>1</sup> Strain abbreviations: Kou – Kounoupidiana, PanA – PanagiaA, Lab – Laboratory.
<sup>2</sup> This gene was also found to be up-regulated in a previous work (Pavlidis et al. 2018) in which it was named contig02103.
<sup>3</sup> This gene is a recently duplicated (paralog) of the previous gene.
<sup>4</sup> This gene was also found to be up-regulated in a previous work (Pavlidis et al. 2018) in which it was named conti00436.

**Table S6.** Number of RNAseq reads (million reads) generated for each sample, for the Malpighian tubule data set.

|  | Replicate 1 | Replicate 2 | Replicate 3 | Total |
| --- | --- | --- | --- | --- |
| **PanagiaB** | 62.74 | 63.62 | 62.82 | 189.18 |
| **Hybrid** | 62.62 | 63.64 | 62.92 | 189.18 |

**Table S8.** List of significantly up-regulated detoxification genes in the Malpighian tubule samples.

| *Gene id* | *log<sub>2</sub>FC* | *FDR* | *Annotation* |
| --- | --- | --- | --- |
| *LOC106621406* | 6.55 | 2.4E-62 | CYP4P6 |
| *LOC106618199* | 3.99 | 6.6E-18 | CYP6G28 |
| *LOC106618206-00001* | 3.91 | 7.4E-04 | CYP6G24 |
| *LOC106618206-00002* | 3.33 | 2.4E-11 | CYP6G6 |
| *LOC106620249* | 2.15 | 8.3E-22 | CYP6A70 |
| *LOC106615161* | 2.36 | 3.57E-23 | GST 1-1 |
| *LOC118679823* | 7.71 | 1.38E-37 | UGT5-like |
| *LOC106623379* | 4.95 | 8.96E-32 | UGT5-like |
| *LOC118682404* | 2.46 | 1.14E-23 | UGT2-like |

**Table S9.**
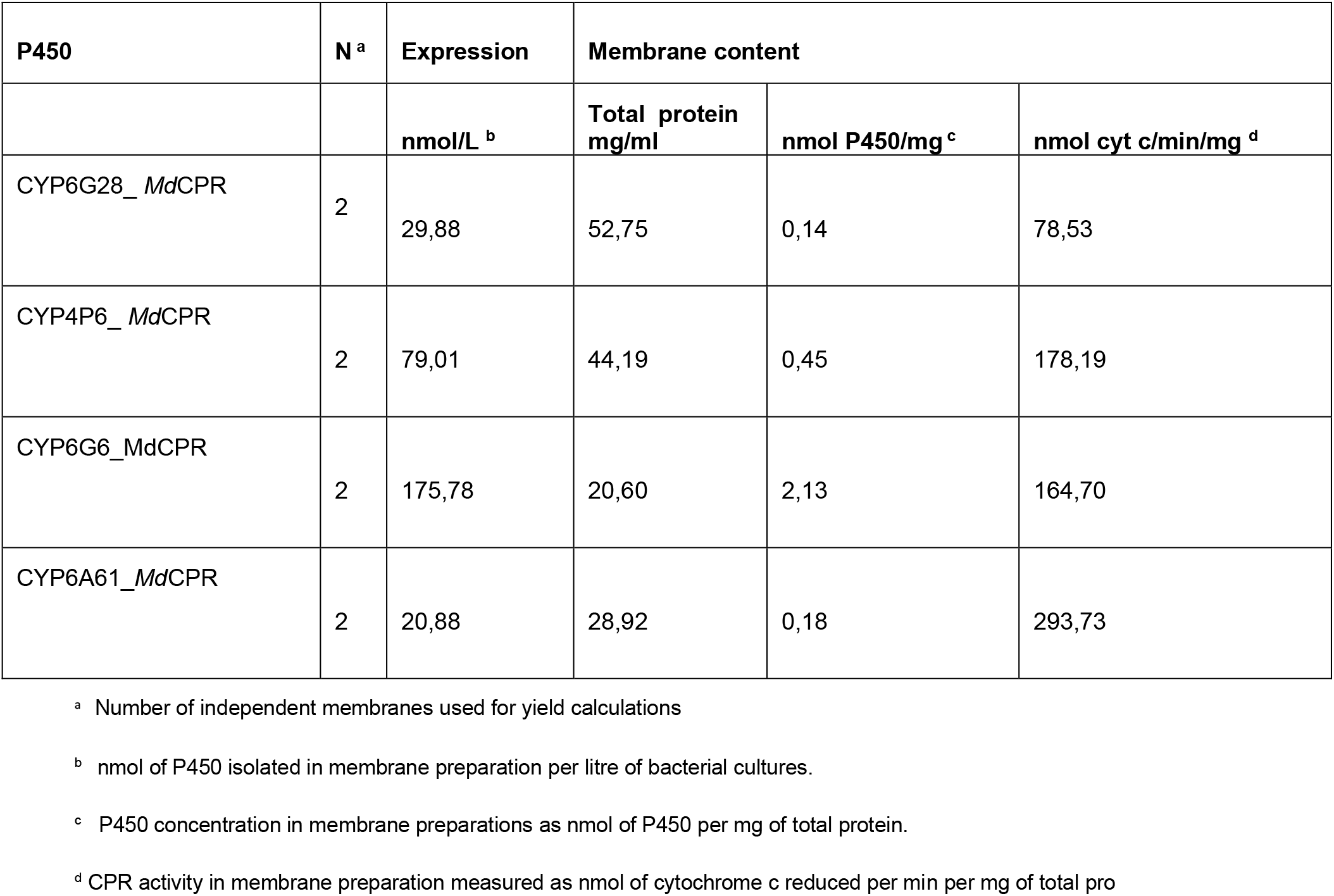
Yields of B. oleae P450 expressed in E. coli

### Figures

**Figure S1.**
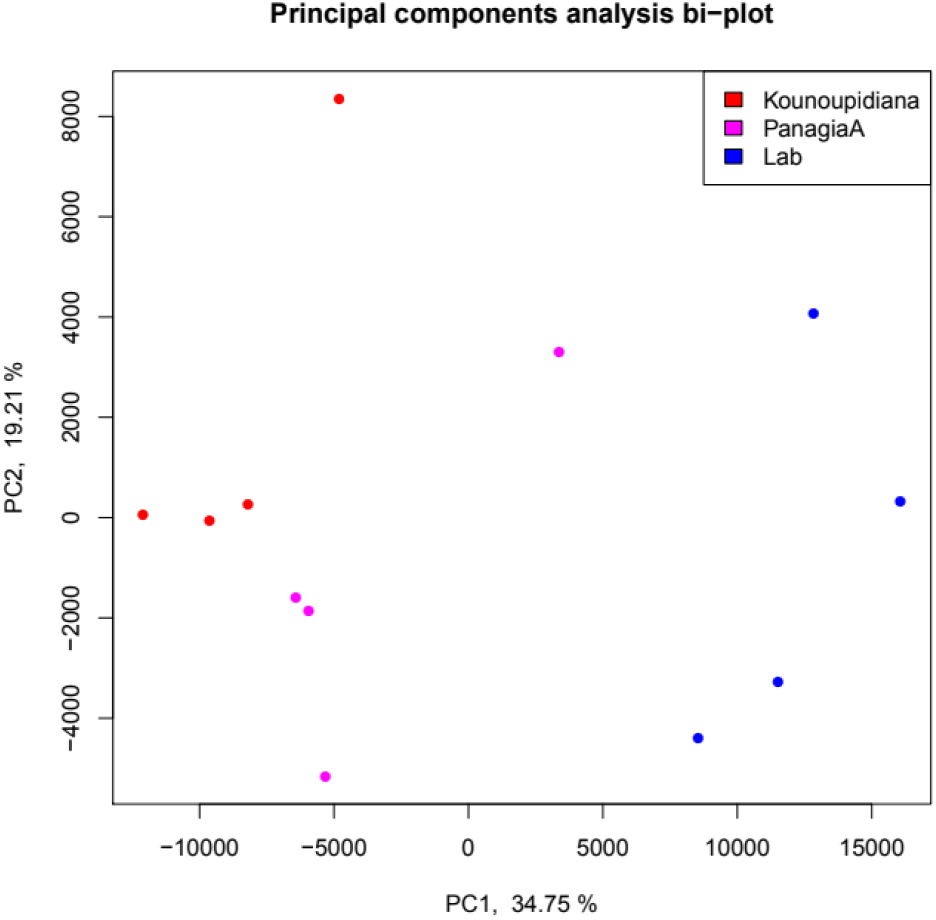
Principal components analysis (PCA) for the whole-body samples.

**Figure S2.**
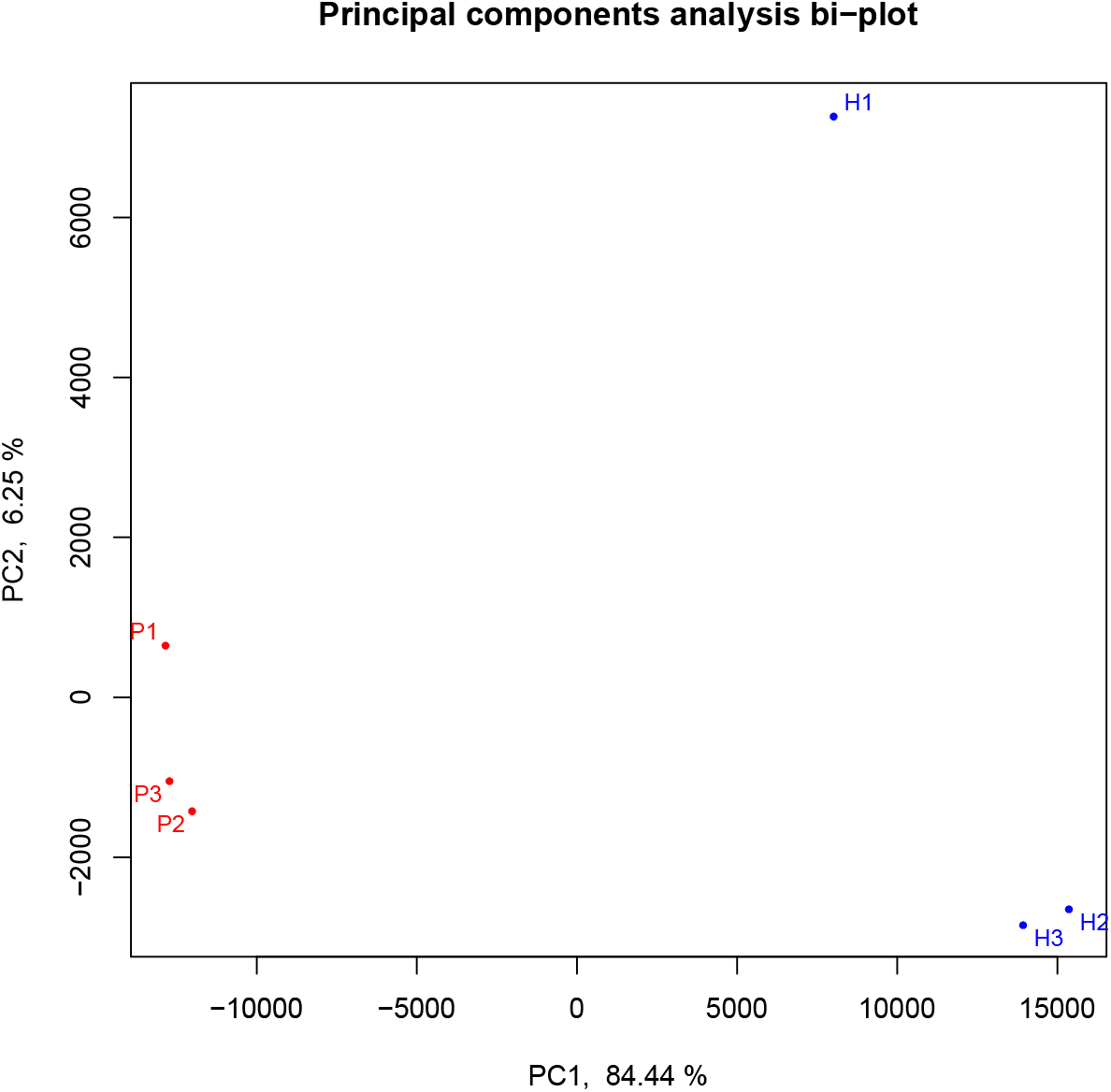
Principal components analysis) for the malpighian tubule samples.

**Figure S3:**
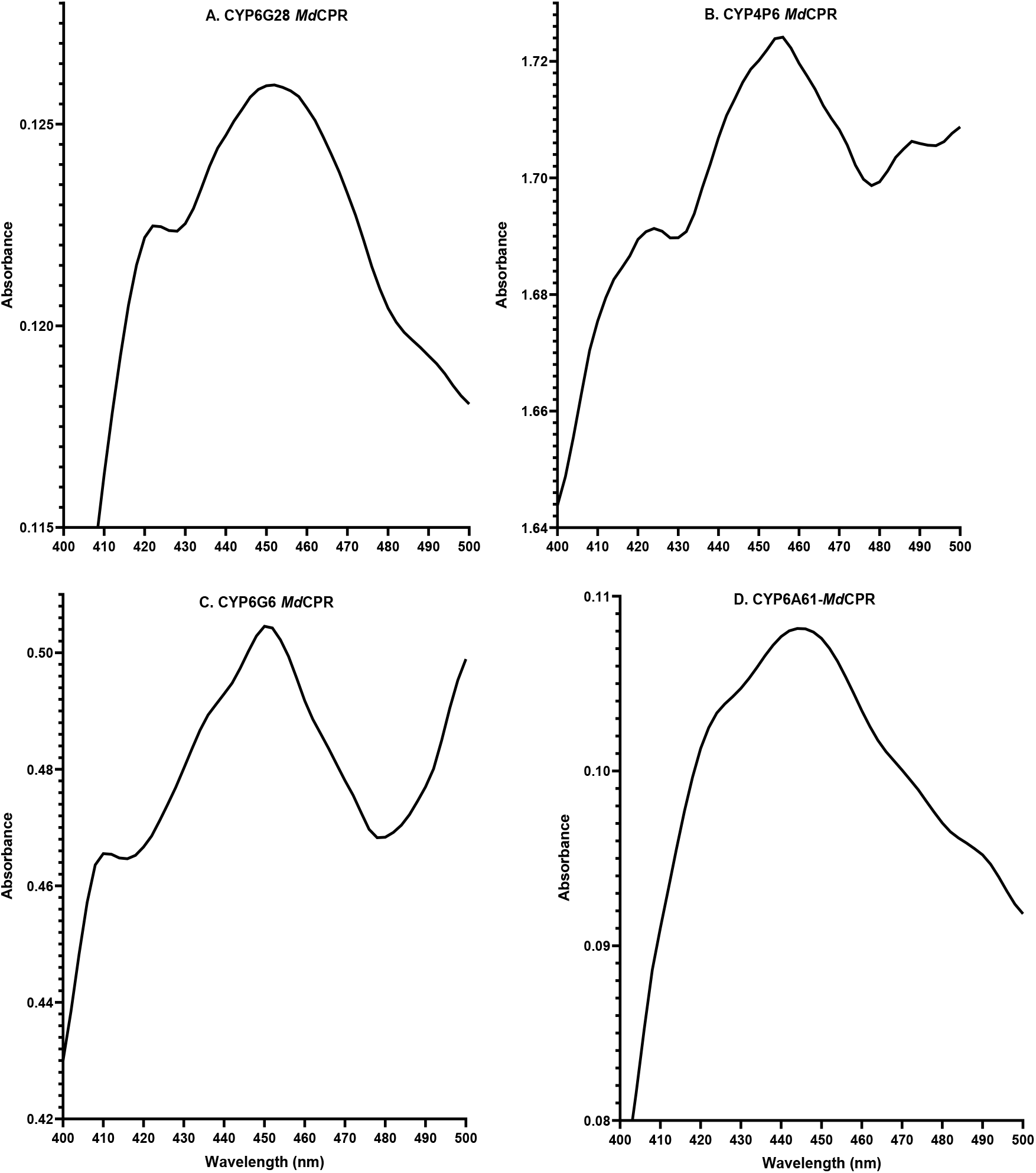
Carbon monoxide difference spectra of bacterial membranes expressing *B. oleae* P450s BoCYP6A61, BoCYP6G6, BoCYP4P6 and BoCYP6G28

